# Mitochondrial Calcium Uptake Promotes Mitochondrial Activation and Hyperactivated Motility During Human Sperm Capacitation

**DOI:** 10.64898/2026.09.28.755036

**Authors:** Pilar Irigoyen, Juan J. Ferreira, Adriana Cassina, Shweta Bhagwat, Nikita Peramsetty, Rossana Sapiro, Celia M. Santi

**Affiliations:** Departamento de Histología y Embriología, Facultad de Medicina, Universidad de la República, Montevideo, Uruguay; Centro de Investigaciones Biomédicas (CEINBIO), Facultad de Medicina, Universidad de la República, Montevideo, Uruguay; Department of Obstetrics and Gynecology, Center for Reproductive Health Sciences, Washington University School of Medicine in St. Louis, St. Louis, MO, United States; Departamento de Bioquímica, Facultad de Medicina, Universidad de la República, Montevideo, Uruguay; Department of Neuroscience, Washington University School of Medicine in St. Louis, St. Louis, MO, United States

**Keywords:** spermatozoa capacitation, Mitochondrial Ca^2+^, CatSper, Mitochondrial calcium uniporter (MCU), Hyperactivated motility

## Abstract

Capacitation is a post-ejaculatory maturation process by which sperm become hyperactive and undergo the acrosome reaction, both of which are required to fertilize an oocyte. Two key features of capacitation are increased intracellular calcium via the calcium channel CatSper and changes in mitochondrial function. In mouse spermatozoa, CatSper-dependent calcium entry promotes mitochondrial calcium uptake and mitochondrial activation, contributing to hyperactivated motility and fertilization ability. Whether this mechanism is conserved in human spermatozoa was unknown. Here, we assessed mitochondrial function and mitochondrial calcium in paired non-capacitated and capacitated human spermatozoa from normozoospermic donors. Capacitation increased mitochondrial respiratory control ratio, coupling efficiency, mitochondrial membrane potential, and mitochondrial calcium. The increase in mitochondrial calcium required extracellular calcium and was prevented by pharmacological inhibition of CatSper or the mitochondrial calcium uniporter (MCU). Inhibition of CatSper or mitochondrial calcium uniporter also attenuated the capacitation-associated increase in mitochondrial membrane potential and reduced hyperactivated motility without affecting progressive or total motility. Pharmacological disruption of mitochondrial function reduced capacitation-associated hyperactivated motility without affecting the acrosome reaction. Together, these findings support a conserved pathway in human and mouse spermatozoa in which CatSper-mediated calcium entry promotes MCU-dependent mitochondrial calcium uptake and mitochondrial activation, contributing to the acquisition of hyperactivated motility during capacitation. These findings identify mitochondrial calcium handling as a potential target for modulating human spermatozoa function.

## Introduction

Spermatozoa acquire fertilization competence through capacitation, which occurs within the female reproductive tract and leads to hyperactivated motility and the acrosome reaction. These responses enable spermatozoa to interact with the female reproductive tract, penetrate the zona pellucida, and fertilize the egg ^1–8^Capacitation in the female reproductive tract is initiated by exposure of spermatozoa to an increased concentration of bicarbonate, which activates soluble adenylyl cyclase, leading to increased cyclic AMP production, protein kinase A activation, and protein phosphorylation ^6,7^. As capacitation progresses, these signaling events are accompanied by membrane hyperpolarization and increased intracellular calcium ion (Ca^2+^) concentration.

In human and mouse spermatozoa, extracellular Ca^2+^ entry is primarily mediated by CatSper, a sperm-specific Ca^2+^ channel localized to the principal piece of the flagellum ^9–11^. Loss of CatSper function causes male infertility in both humans and mice, as CatSper-deficient spermatozoa fail to develop normal hyperactivated motility ^12–16^. However, the full range of mechanisms through which CatSper-dependent Ca^2+^ signaling regulates hyperactivation continues to be defined.

One potential downstream target of CatSper-dependent Ca^2+^ signaling is mitochondria. Hyperactivated motility imposes substantial energetic demands ^17,18^, while mitochondrial activity increases during human sperm capacitation ^19,20^. These observations have raised interest in understanding the contribution of mitochondrial function to the energetic demands of sperm capacitation. Beyond their role in energy metabolism production, mitochondria are also Ca^2+^-responsive organelles. In many cell types, increases in mitochondrial Ca^2+^ regulate Ca^2+^-sensitive enzymes involved in oxidative metabolism ( Kumar, 2023; Rossi et al., 2019a) In mouse spermatozoa, mitochondrial Ca^2+^ increases during capacitation via a mechanism dependent on extracellular Ca^2+^ and CatSper activity. Additionally, mitochondrial Ca^2+^ uptake through the mitochondrial calcium uniporter (MCU) contributes to mitochondrial function, hyperactivated motility, and fertilization ^21,23–26^. Whether this mechanism is conserved in human spermatozoa is unknown.

It may not be possible to directly extrapolate findings from mouse sperm to human sperm given species-specific differences in CatSper regulation and spermatozoa energy metabolism. For example, CatSper activation and regulation differ between human and mouse sperm, including differences in their sensitivity to specific channel modulators ^27–34^. Additionally, the relative contribution and regulation of glycolysis and oxidative phosphorylation can differ between species (Vicente-Carrillo et al., 2023). Moreover, defects in human CatSper function can cause infertility despite normal semen parameters ^16,31,35–37^. We hypothesized that CatSper-mediated Ca^2+^ entry during human spermatozoa capacitation promotes MCU-dependent mitochondrial Ca^2+^ uptake and mitochondrial activation, thereby contributing to the acquisition of hyperactivated motility. To test this hypothesis, we assessed mitochondrial respiration, mitochondrial membrane potential, NADH autofluorescence, and mitochondrial Ca^2+^ in paired non-capacitated and capacitated human spermatozoa. We further disrupted mitochondrial function and pharmacologically inhibited CatSper and MCU to determine the contribution of mitochondrial activity and Ca^2+^ handling to hyperactivated motility and the acrosome reaction.

### Materials and Methods

#### Reagents

All reagents used to prepare non-commercial Biggers–Whitten–Whittingham (BWW) media were obtained from Millipore Sigma/Sigma-Aldrich (St. Louis, MO, USA) unless otherwise specified. Calcium ionophore A23187 (catalog #52665-69-7), dimethyl sulfoxide (DMSO; D8418), sodium cyanide (NaCN; 205222; ACS reagent, ≥95%), antimycin A (AA; A8674), carbonyl cyanide 4-(trifluoromethoxy) phenylhydrazone (FCCP; CAS 370-86-5), Ru360 (557440; ≥97%), and propidium iodide (PI; 537059; CAS 25535-16-4) were obtained from Millipore Sigma/Sigma-Aldrich. Oligomycin (HY-N6782; CAS 1404-19-9), NNC 55-0396 dihydrochloride (HY-50722), and Calcein-AM (HY-D0041; CAS 148504-34-1) were obtained from MedChemExpress (Monmouth Junction, New Jersey, USA). APC-conjugated mouse anti-human CD46 monoclonal antibody (C-10; sc-166159) was obtained from Santa Cruz Biotechnology (Dallas, TX, USA). Fluo-4 AM (F14201/F14204), Fluo-5N AM, MitoTracker Red CMXRos (M22426), Pluronic F-127 (P6867), and tetramethylrhodamine methyl ester perchlorate (TMRM; T668) were obtained from Invitrogen/Thermo Fisher Scientific (Waltham, MA, USA).

#### Subject Selection and Ethical Approval

Study protocols were approved by the Ethics Committee of the Facultad de Medicina, Universidad de la República (UDELAR, Montevideo, Uruguay) and by the Washington University in St. Louis (WashU) Institutional Review Board (IRB ID #202507087). Written informed consent was obtained from all participants before sample collection. Participant anonymity was maintained throughout the study, and researchers had no access to identifying information.

The study included men attending the Fertilab-Reprovita andrology clinics (Montevideo, Uruguay) for routine semen analysis between March 2022 and May 2026, as well as volunteer donors recruited through the WashU Fertility & Reproductive Medicine Center between September 2022 and July 2026. Freshly ejaculated semen samples were collected by masturbation after 3–7 days of abstinence in accordance with the World Health Organization Laboratory Manual for the Examination and Processing of Human Semen ^38^. Samples were collected into sterile plastic containers and processed after liquefaction. Samples were included if the donors were classified as normozoospermic according to WHO (2021) reference criteria, irrespective of participant fertility status. Diagnostic semen analyses were performed at the Fertilab-Reprovita laboratories or the WashU Fertility & Reproductive Medicine Center. Except for experiments assessing Ca^2+^ dynamics, all experiments followed a paired design in which aliquots from the same semen sample were distributed among the experimental conditions for a given assay, allowing each donor to serve as their own control. Donors were allocated to individual experiments according to sample and biological material availability, and sample sizes varied among assays.

#### Semen Evaluation and Computer-Assisted Spermatozoa Analysis

Fresh semen samples were allowed to liquefy at room temperature for 30 min, and then semen parameters were evaluated according to WHO guidelines ^38^. Hyperactivated motility was assessed among motile spermatozoa according to three computer-assisted semen analysis (CASA)-derived kinematic parameters: curvilinear velocity (VCL), linearity (LIN), and amplitude of lateral head displacement (ALH), according to established approaches ^39–41^. Because CASA measurements can vary depending on the acquisition and analysis system, laboratory-specific thresholds were used and optimized for each system. Hyperactivation was expressed as the percentage of motile spermatozoa meeting all three corresponding criteria.

At UDELAR, sperm concentration and motility parameters were assessed with a CASA system (SCA Evolution; Microptic, Barcelona, Spain). Semen aliquots were loaded into two prewarmed CELL-VU sperm counting chambers (DRM-600; Millennium Sciences, Inc., New York, NY, USA), and 10 randomly selected fields per chamber were analyzed on a Nikon Eclipse E200 microscope equipped with a Basler acA780-75gc camera at 100× magnification and maintained at 37 °C. Spermatozoa were classified as hyperactivated if VCL was >150 µm/s, LIN was <50%, and ALH was >3.5 µm. A minimum of 200 spermatozoa were analyzed per sample.

At WashU, sperm concentration and motility parameters were assessed by loading 3 µL of sample into a prewarmed (37 °C), 20-µm-deep Leja Standard Count 4 chamber. CASA was performed on a Hamilton-Thorne CEROS II digital image analyzer (v1.7; Hamilton-Thorne Research, Beverly, MA, USA). Spermatozoa were classified as hyperactivated if VCL was >125 µm/s, LIN was <50%, and ALH was >5.0 µm. A minimum of 200 spermatozoa were analyzed per sample.

#### Spermatozoa Preparation and Capacitation

Liquefied semen (500 µL) was subjected to swim-up in 1 mL BWW medium containing 95 mM NaCl, 4.6 mM KCl, 1.7 mM CaCl_2_, 1.2 mM KH_2_PO_4_, 1.2 mM MgSO_4_, 5.6 mM glucose, 0.27 mM sodium pyruvate, 20 mM HEPES, and 44 mM sodium lactate [26]. Samples were incubated for 1 h at 37 °C with tubes inclined at 45°, after which 500 µL of the upper fraction was collected.

Motile spermatozoa were divided into two aliquots. One aliquot was maintained under non-capacitating conditions (NC), whereas the other was supplemented with 25 mM NaHCO_3_ and 5 mg/ml BSA to induce capacitation (CAP). Where indicated, solutions additionally contained 2.5 µM antimycin A (AA), 5 µM FCCP, 20 µM Ru360, 1 µM NNC 55-0396, or 30 µM progesterone. NC and CAP aliquots were incubated in parallel for 4 h at 37 °C before analysis.

For high-resolution respirometry experiments, semen samples were processed differently because of the larger number of spermatozoa required. Liquefied semen was centrifuged at 500 × *g* for 10 min at room temperature to separate spermatozoa from seminal plasma. The spermatozoa pellet was subsequently resuspended in either NC or CAP BWW medium and incubated under the same conditions described above.

#### Mitochondrial Function by High-Resolution Respirometry

Oxygen consumption rate was measured in 24 × 10^6^ spermatozoa under paired NC and CAP conditions in an Oxygraph-2k high-resolution respirometer (Oroboros Instruments GmbH, Innsbruck, Austria) as described ^42^. Mitochondrial respiration was assessed in intact spermatozoa suspended in BWW medium. Basal respiration was recorded before sequential addition of 2 µg/mL oligomycin to inhibit ATP synthase, FCCP (0.1–1 µM) to achieve maximal uncoupled respiration, and 2.5 µM AA to inhibit mitochondrial respiratory complex III. Residual oxygen consumption following AA treatment was considered non-mitochondrial respiration and was subtracted from the other respiratory measurements before parameter calculation. Oxygen consumption rate values were normalized to 10^6^ spermatozoa. Respiratory control ratio (RCR), coupling efficiency, and spare respiratory capacity were calculated as follows: RCR = maximal respiration/proton leak; coupling efficiency = (basal respiration − proton leak)/basal respiration; and spare respiratory capacity = maximal respiration/basal respiration. Because these indices represent ratios of respiratory parameters, they are independent of spermatozoa number.

#### Flow Cytometric Assessment of Mitochondrial Membrane Potential, Ca^2+^, and Acrosome Reaction

Spermatozoa concentration was adjusted to 1–4 × 10^6^ cells/mL. Depending on the assay, samples were incubated at 37 °C with 300 nM TMRM to assess mitochondrial membrane potential (MMP), 0.2 µg/µL APC-conjugated mouse anti-human CD46 monoclonal antibody (C-10; sc-166159, Santa Cruz Biotechnology) to assess the acrsosome reaction, 2–4 µM Fluo-5N AM supplemented with 0.05% Pluronic F-127 for relative mitochondrial Ca^2+^ measurements, or Fluo-4 AM for cytosolic Ca^2+^ measurements. TMRM, CD46 antibody, Fluo-5N AM, and Fluo-4 AM were added 25, 60, 90, and 45 min, respectively, before completion of the capacitation period.

Cell viability was assessed by incubating samples for 15 min at 37 °C in either 0.1 µM Calcein-AM or 1 µg/mL propidium iodide. After staining, excess probe was removed by centrifugation at 500 × *g* for 10 min, and spermatozoa were resuspended in 300 µL PBS or NC BWW medium.

For MMP measurements, each washed sample was divided into two aliquots. One aliquot was analyzed under the corresponding experimental condition, whereas the second was treated with 20 µM FCCP for 15 min to induce mitochondrial depolarization and provide a minimum-fluorescence (Fmin) reference for TMRM normalization.

For mitochondrial Ca^2+^ measurements, each sample was divided into three aliquots: one representing the experimental condition, a second treated with 2–5 µM ionomycin to determine maximal fluorescence (Fmax), and a third maintained in Ca^2+^-free medium supplemented with 2 mM EGTA to determine Fmin. These reference conditions were used for fluorescence normalization and to minimize variability arising from differences in probe loading. For acrosome reaction assessment, spermatozoa were stimulated with 10 µM A23187 before analysis.

Flow cytometric analysis of TMRM fluorescence was performed either at UDELAR by on a BD Accuri C6 Plus flow cytometer (BD Biosciences, San Jose, CA, USA) or at WashU on a Cytek Aurora spectral flow cytometer (Cytek Biosciences, Fremont, CA, USA). All other flow cytometry experiments were performed at WashU on the Cytek Aurora. Identical staining protocols, gating strategies, and data-analysis criteria were applied to TMRM experiments performed at both sites. Forward-scatter (FSC-H) and side-scatter (SSC-H) characteristics were used to exclude debris. Single cells were selected according to FSC-H versus FSC-A gating to exclude doublets. At least 25,000 single-cell events were acquired per sample. BD Accuri C6 Plus software or SpectroFlo software, as appropriate, were used for data acquisition and analysis. For spectral flow cytometry experiments, unstained and single-stained controls were included to generate the spectral unmixing matrix.

#### NADH Autofluorescence Measurements

At the end of the 4-h incubation, paired NC and CAP samples were plated at 2 × 10^6^ cells in a final volume of 200 µL per well in a 96-well plate and analyzed using a BioTek Synergy H1 multimode plate reader (BioTek Instruments). Fluorescence was measured in a top-read configuration.

NAD(P)H autofluorescence, representing the combined fluorescence of reduced NADH and NADPH, was measured as an indicator of cellular redox state. Because NADH and NADPH have overlapping fluorescence spectra, this approach does not distinguish their individual contributions to the measured signal. Furthermore, changes in NAD(P)H autofluorescence reflect the balance between NAD(P)H production and oxidation and therefore cannot, by themselves, distinguish changes in production from changes in oxidation.

NAD(P)H autofluorescence was spectrally characterized in human spermatozoa to optimize the excitation and emission wavelengths used for subsequent measurements. Based on this optimization, an excitation wavelength of 310 nm was selected. Emission spectra were then acquired following excitation at 310 nm in NC and CAP spermatozoa under control conditions and following FCCP treatment **(Supplementary Figure 1)**. Based on these spectral measurements, NAD(P)H autofluorescence was quantified using excitation at 310 nm and emission at 450 nm.

For kinetic measurements, NAD(P)H autofluorescence was recorded every minute for 10 min under basal conditions. FCCP (20 µM) or NaCN (100 µM) was then added, and fluorescence was recorded every minute for an additional 10 min. Paired NC and CAP samples from the same donor were analyzed in parallel.

For the experiments presented in Figure 2, NAD(P)H autofluorescence in NC and CAP spermatozoa was normalized independently to the Fmin obtained following FCCP treatment of the corresponding sample. Thus, the FCCP response within each condition served as its respective Fmin reference. In a subset of experiments used to independently validate this normalization approach **(Supplementary Figure 1)**, fluorescence was normalized using FCCP-induced fluorescence as Fmin and NaCN-induced fluorescence as Fmax. This approach provided a defined fluorescence range for comparison of NAD(P)H autofluorescence between paired NC and CAP spermatozoa.

#### Colocalization of Fluo-5N with Mitochondria

Fluo-5N loading and mitochondrial colocalization imaging were performed as described for mouse spermatozoa [15], with modifications for human spermatozoa. Spermatozoa were loaded with 2–4 µM Fluo-5N AM in BWW medium containing 0.05–0.1% Pluronic F-127 for 90 min at 37 °C. MitoTracker Red CMXRos (200 nM) was added during the final 20–30 min of incubation. After staining, spermatozoa were washed and allowed to adhere to coverslips coated with 0.1% poly-L-lysine. Images were acquired at 22–23 °C on a Zeiss LSM 980 Airyscan 2 microscope with a 100× objective at a resolution of 2048 × 2048 pixels. Fluo-5N and MitoTracker were excited at 488 and 543 nm, respectively, and fluorescence emission was collected at 500–530 nm and >560 nm. Colocalization was analyzed with ImageJ v1.48.

#### Statistical Analysis

GraphPad Prism version 8.0.1 for Windows (GraphPad Software, San Diego, CA, USA) was used for all statistical analyses. Data are presented as mean ± standard deviation unless otherwise indicated. Paired or unpaired tests were performed according to the experimental design and as specified in the corresponding figure legends. For comparisons between paired NC and CAP spermatozoa, paired *t*-tests were performed. In experiments involving independent groups or additional pharmacological conditions, the statistical test performed is specified in the corresponding figure legend. Statistical significance was defined as *p* < 0.05.

Sample size (*n*) represents the number of independent donors unless otherwise specified. Sample sizes varied among experiments because the number of assays that could be performed from an individual semen sample depended on the amount of biological material available. Technical replicates were not considered independent biological replicates.

## Results

### Mitochondrial function increases during human spermatozoa capacitation

To characterize changes in mitochondrial function during human spermatozoa capacitation, we first assessed mitochondrial respiration by high-resolution respirometry in paired non-capacitated (NC) and capacitated (CAP) spermatozoa **(Figure 1; Table 1)** ^42,43^. CAP spermatozoa exhibited significantly higher respiratory control ratio (RCR) **(Figure 1A; Table 1)**, coupling efficiency **(Figure 1B; Table 1)**, basal respiration, proton leak, and maximum respiratory rate than paired NC spermatozoa **(Table 1)**. In contrast, spare respiratory capacity was similar between CAP and NC spermatozoa **(Figure 1C; Table 1)**. Together, these findings indicate enhanced mitochondrial respiratory function after capacitation, consistent with previous reports in human spermatozoa ^20^.

**Figure 1.**
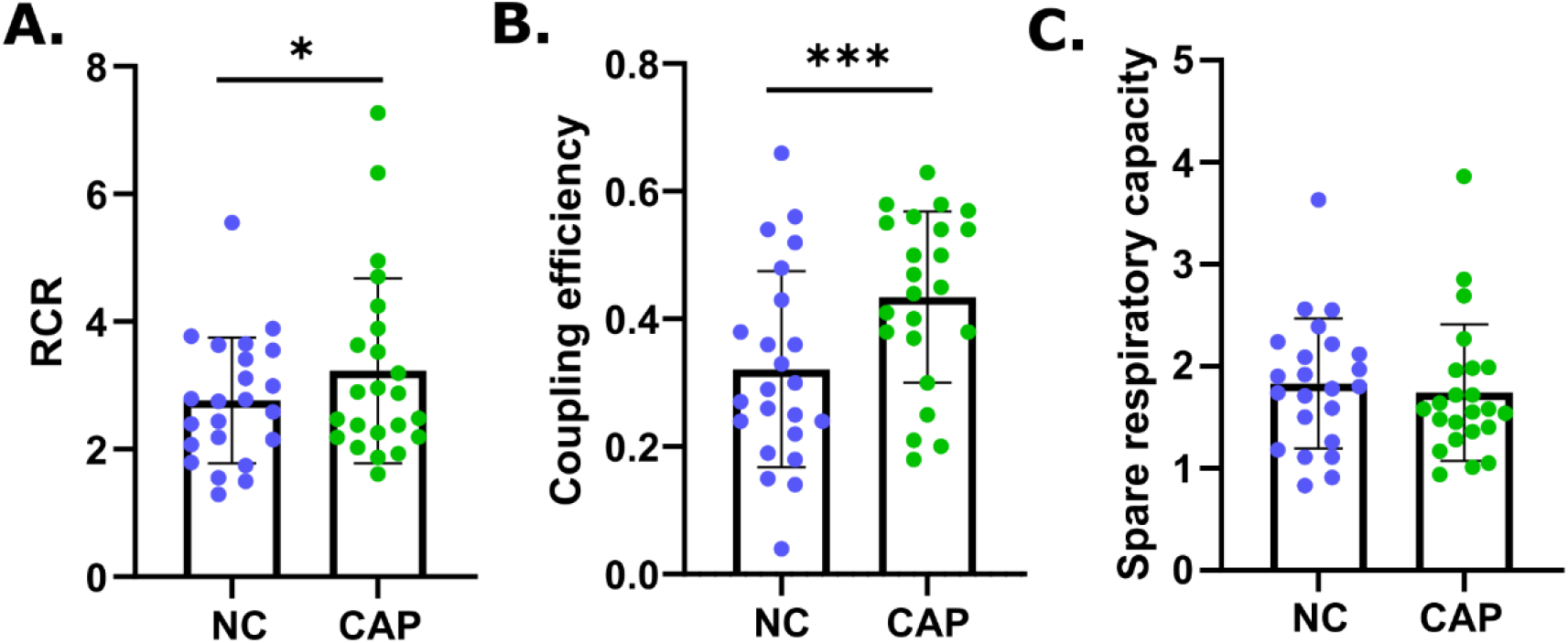
Mitochondrial respiratory function increases during human spermatozoa capacitation. Mitochondrial respiratory indices calculated from high-resolution respirometry measurements of intact non-capacitated **(NC; blue)** and capacitated **(CAP; green)** human spermatozoa. **(A)** Respiratory control ratio **(RCR)**, calculated as maximal respiration divided by proton leak. **(B)** Coupling efficiency, calculated as (basal respiration − proton leak)/basal respiration. **(C)** Spare respiratory capacity, calculated as maximal respiration divided by basal respiration. * *P* < 0.05; *** *P* < 0.001 by paired t-test; n = 23.

**Table 1.** Respiratory parameters and indices derived from high-resolution respirometry in non-capacitated (NC) and capacitated (CAP) human spermatozoa. Basal respiration represents oxygen consumption in the presence of exogenous substrates, whereas proton leak corresponds to oligomycin-resistant respiration. Maximal respiration was determined following dissipation of the mitochondrial proton gradient with FCCP, and non-mitochondrial respiration was defined as residual oxygen consumption after inhibition of mitochondrial respiration with antimycin A. Respiratory indices were calculated from these parameters as follows: respiratory control ratio (RCR) = maximal respiration/proton leak; coupling efficiency = (basal respiration − proton leak)/basal respiration; and spare respiratory capacity = maximal respiration/basal respiration. Data are presented as *n*, mean, minimum (Min), maximum (Max), and standard deviation (SD). *n*, number of donors. A paired *t*-test was used for statistical comparisons between paired NC and CAP samples.

|  | n | Mean |  | Min |  | Max |  | SD |  | p-value |
| --- | --- | --- | --- | --- | --- | --- | --- | --- | --- | --- |
| Parameters (pmol O<sub>2</sub> * sec<sup>-1</sup> * mL<sup>-1</sup> * 10<sup>-6</sup> cells) |  | NC | CAP | NC | CAP | NC | CAP | NC | CAP |  |
| Basal respiration | 23 | 0.523 | 0.830 | 0.060 | 0.130 | 1.790 | 2.560 | 0.442 | 0.709 | <0,0001**** |
| Proton leak |  | 0.363 | 0.452 | 0.040 | 0.070 | 1.320 | 1.280 | 0.335 | 0.383 | 0.0433* |
| Maximum respiratory rate |  | 0.977 | 1.303 | 0.060 | 0.130 | 3.230 | 3.700 | 0.894 | 0.961 | 0.0183* |
| Non-mitochondrial respiration |  | 0.392 | 0.511 | 0.030 | 0.040 | 2.070 | 2.500 | 0.475 | 0.613 | 0.008** |
| **Indices** |  |  |  |  |  |  |  |  |  |  |
| RCR |  | 2.763 | 3.227 | 1.290 | 1.610 | 5.550 | 7.270 | 0.988 | 1.452 | 0.0257* |
| Coupling efficiency |  | 0.321 | 0.434 | 0.040 | 0.180 | 0.660 | 0.630 | 0.153 | 0.134 | 0.0003*** |
| Spare respiratory capacity |  | 1.831 | 1.742 | 0.830 | 0.940 | 3.630 | 3.860 | 0.637 | 0.668 | 0.2252 |

We next sought to identify changes in mitochondrial function that underlie the increased respiration observed in CAP spermatozoa. Thus, we used TMRM fluorescence to assess mitochondrial membrane potential, which reflects the electrochemical gradient across the inner mitochondrial membrane generated by respiratory chain activity. We also measured NADH autofluorescence as an indicator of the intracellular pool of reduced NADH and, consequently, cellular redox state. TMRM fluorescence was higher **(Figure 2A)**, and NADH autofluorescence was lower **(Figure 2B)**, in CAP spermatozoa than in NC spermatozoa. The possibility that lower NADH levels reflect a reduced supply of reducing equivalents rather than increased NADH oxidation cannot be excluded. Together, these findings demonstrate that capacitation in human spermatozoa is associated with increased mitochondrial respiration and mitochondrial membrane potential, accompanied by increase in NADH oxidation.

**Figure 2.**
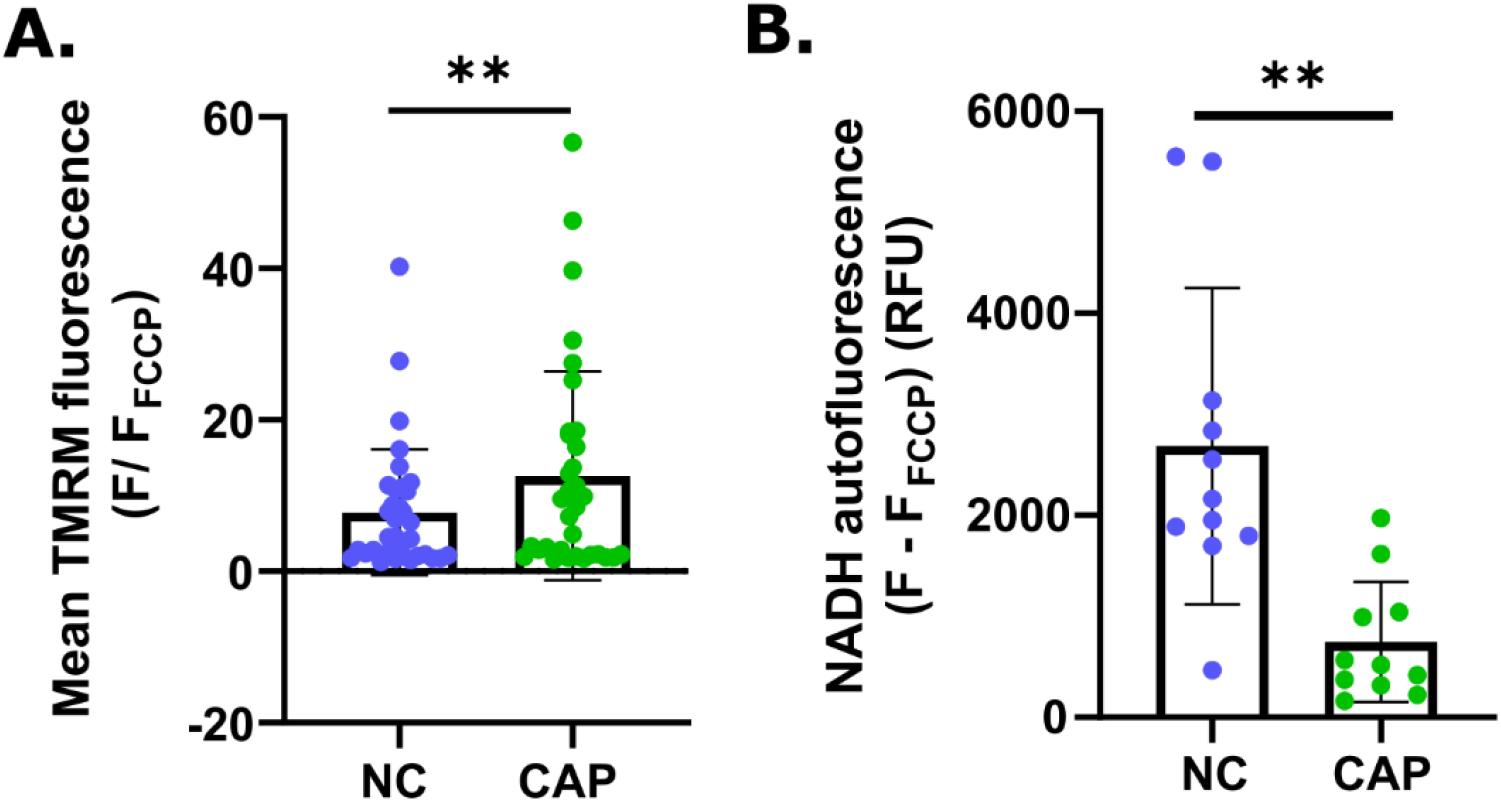
Capacitated human spermatozoa exhibit increased mitochondrial membrane potential and decreased NADH autofluorescence. **(A)** Mitochondrial membrane potential assessed by flow cytometric analysis of TMRM fluorescence (n = 34). TMRM fluorescence was normalized to the minimum fluorescence obtained after FCCP treatment of the same sample. Data were collected in parallel at the Facultad de Medicina, Universidad de la República (n = 13) and Washington University in St. Louis (n = 21). **(B)** NADH autofluorescence measured in a plate reader (λ_ex = 310 nm; λ_em = 450 nm) and normalized to the minimum fluorescence obtained after FCCP treatment (n = 11). RFU, relative fluorescence units; NC, non-capacitated spermatozoa (blue); CAP, capacitated spermatozoa (green). ** *P* < 0.01 by paired t-test.

### Mitochondrial activity contributes to hyperactivation but not the acrosome reaction during human spermatozoa capacitation

In mouse spermatozoa, inhibition of mitochondrial function significantly impairs acquisition of hyperactivated motility but not the acrosome reaction during capacitation ^23^. Thus, we next asked whether mitochondrial activity was required for hyperactivated motility and the acrosome reaction in human spermatozoa. To do so, we disrupted mitochondrial function with AA or FCCP. As expected, the percentage of hyperactivated spermatozoa increased significantly following capacitation (0.425 ± 0.942% vs. 12.750 ± 8.533%) **(Figure 3A)**. Disruption of mitochondrial function significantly reduced this increase, with the percentage of hyperactivated spermatozoa decreasing to 5.330 ± 6.252% in the presence of AA and to 1.613 ± 2.569% in the presence of FCCP (**Figure 3A**). In contrast, progressive motility was not significantly different among conditions (**Supplementary Figure 2**) confirming that cell viability was not affected. Neither AA nor FCCP significantly altered spontaneous or A23187-induced acrosome reaction (**Figure 3B**). Thus, as observed in mouse spermatozoa, disruption of mitochondrial function impaired the acquisition of hyperactivated motility in human spermatozoa without significantly affecting progressive motility or the acrosome reaction.

**Figure 3.**
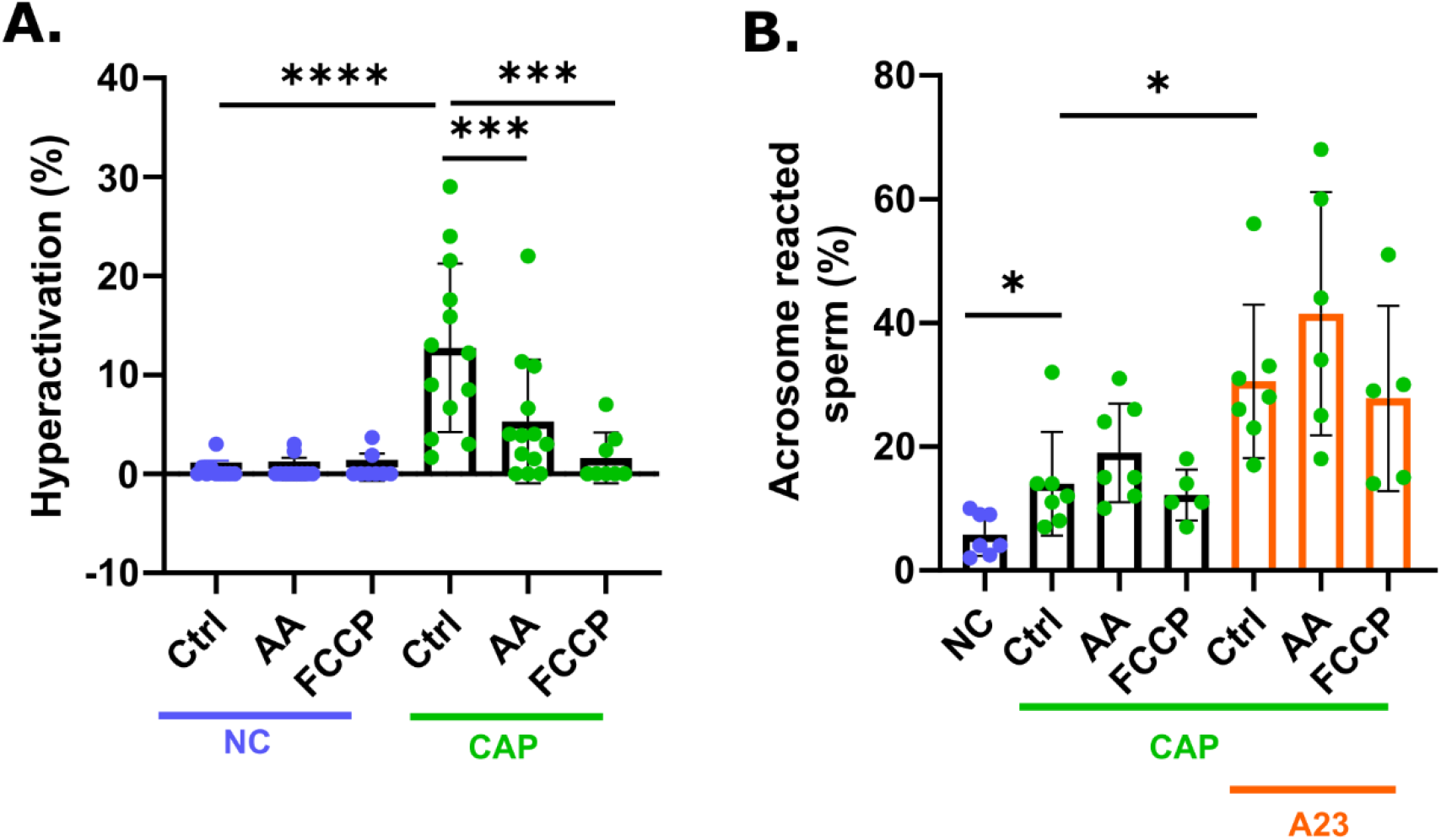
Mitochondrial dysfunction impairs hyperactivation but not the acrosome reaction in human spermatozoa. Spermatozoa were incubated for 4 h under non-capacitating (NC; blue) or capacitating (CAP; green) conditions in the presence of vehicle (Ctrl), 5 µM FCCP, or 2.5 µM antimycin A (AA). **(A)** Percentage of hyperactivated spermatozoa assessed by computer-assisted spermatozoa analysis (CASA) and expressed as a percentage of motile spermatozoa (n = 13). **(B)** Spontaneous and A23187-induced (A23; orange) acrosome reaction, assessed by flow cytometry with an anti-CD46 antibody (n = 5–7). Progressive motility as a vitality control is shown in **Supplementary Figure 2**. * *P* < 0.05; *** *P* < 0.001; **** *P* < 0.0001 by paired t-test.

### Mitochondrial Ca^2+^ increases during human spermatozoa capacitation through a CatSper- and MCU-dependent mechanism

In mouse spermatozoa, mitochondrial Ca^2+^ increases during capacitation, and this increase depends on extracellular Ca^2+^, CatSper activity, and mitochondrial calcium uniporter (MCU) activity ^23^. To determine whether this mitochondrial Ca^2+^ response and its underlying mechanism are conserved in human spermatozoa, we first determined whether the Ca^2+^-sensitive dye Fluo-5N could be used to assess mitochondrial Ca^2+^ in human spermatozoa. Fluo-5N fluorescence was concentrated in the spermatozoa midpiece, where the mitochondrial sheath is located **(Figure 4A,C)**. Analysis of fluorescence intensity across the midpiece revealed the characteristic profile expected from the mitochondrial sheath, with decreased fluorescence in the axoneme region **(Figure 4C,F)**. Fluo-5N fluorescence closely overlapped with the mitochondrial marker MitoTracker Red **(Figure 4B–G)**, supporting the localization of Fluo-5N to the mitochondrial compartment in human spermatozoa.

**Figure 4.**
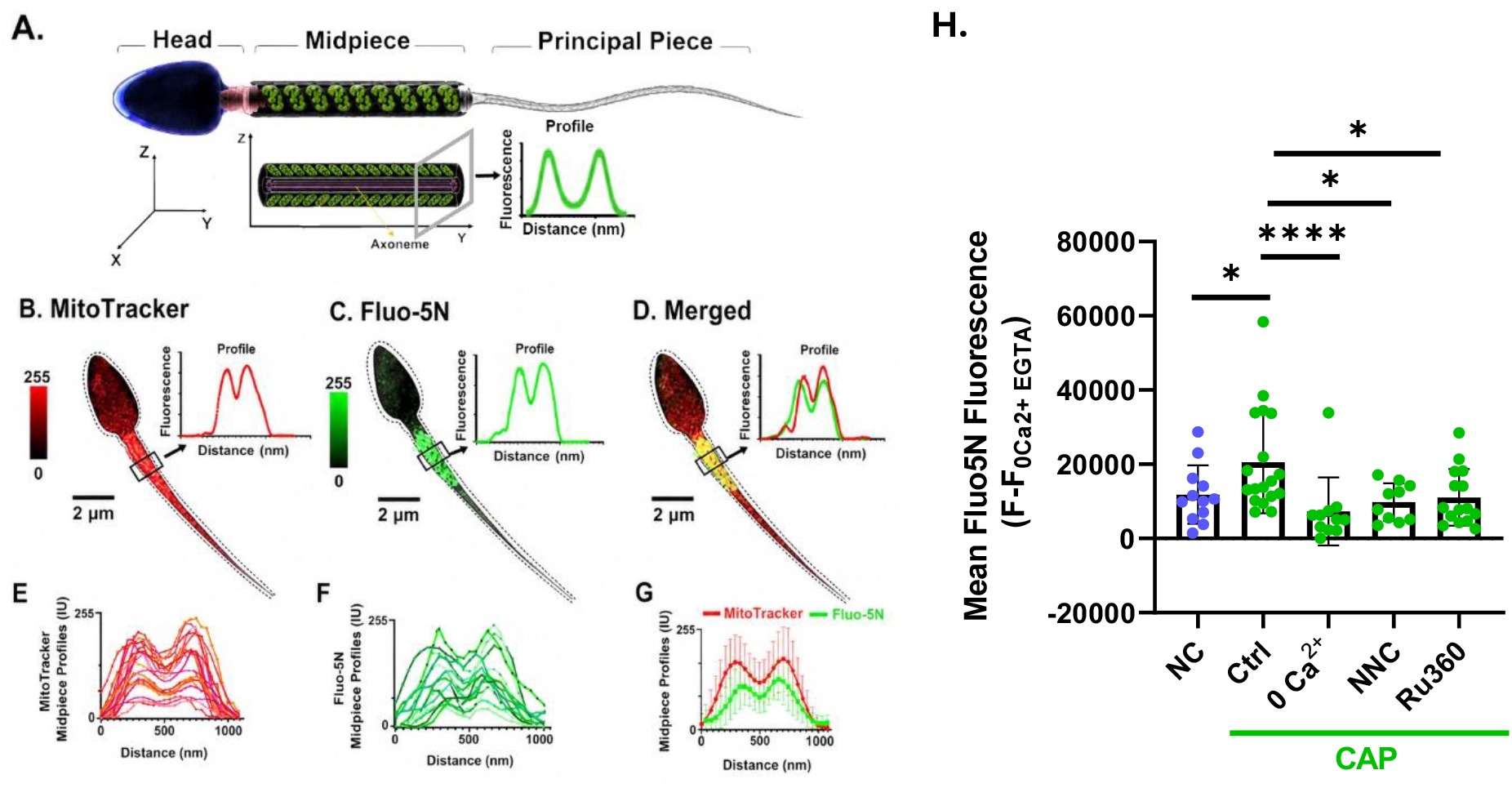
Fluo-5N localizes to the mitochondrial compartment, and mitochondrial Ca^2+^ increases during human spermatozoa capacitation through a CatSper- and MCU-dependent mechanism. **(A)** Schematic representation of human spermatozoa morphology showing the mitochondrial sheath surrounding the axoneme in the midpiece. Cross-sectional and longitudinal representations illustrate the organization of the mitochondrial sheath and the optical plane used to analyze fluorescence intensity across the midpiece. **(B, C)** Representative human spermatozoon labeled with (**B**) MitoTracker Red and (**C**) Fluo-5N and corresponding fluorescence intensity profiles across the midpiece. **(D)** Merged MitoTracker Red and Fluo-5N signals and corresponding fluorescence profiles. **(E)** MitoTracker Red and (**F**) Fluo-5N fluorescence intensity profiles across the midpiece of individual spermatozoa. A total of 15 spermatozoa from 3 donors were analyzed in E and F. **(G)** Mean MitoTracker Red and Fluo-5N fluorescence profiles across the spermatozoa midpiece. Data are presented as mean ± SD. **(H)** Relative mitochondrial Ca^2+^ concentrations measured by Fluo-5N fluorescence in non-capacitated (NC; blue; n = 12) and capacitated (CAP; green; n = 12) spermatozoa in control (ctrl; n=19) conditions, Ca^2+^-free medium (0 Ca^2+^; n = 11), in the presence of 1 µM NNC 55-0396 (NNC; n = 10), or in the presence of 20 µM Ru360 (n = 15). Fluo-5N fluorescence was normalized to the minimum fluorescence obtained for each sample under Ca^2+^-free conditions in the presence of EGTA. Each symbol in H represents an individual donor. Flow cytometry measurements were performed at Washington University in St. Louis. \**P* < 0.05; **** *P* < 0.0001 by independent t-tests.

We next asked whether mitochondrial Ca^2+^ increases during human spermatozoa capacitation. Relative mitochondrial Ca^2+^ concentrations were significantly higher in CAP than in paired NC spermatozoa (20,544 ± 13,790 vs. 11,810 ± 7,861) **(Figure 4H)**. This capacitation-associated increase was abolished when spermatozoa were capacitated in Ca^2+^-free medium (7,287 ± 9,171) **(Figure 4H)**, indicating that the increase in mitochondrial Ca^2+^ requires extracellular Ca^2+^. To determine whether CatSper, the major Ca^2+^ channel activated during capacitation, mediates this increase in mitochondrial Ca^2+^, we treated spermatozoa with the CatSper inhibitor NNC 55-0396. This prevented the capacitation-associated increase in mitochondrial Ca^2+^ (9,800 ± 5,105) **(Figure 4H)**.

To determine whether the capacitation-associated increase in mitochondrial Ca^2+^ requires MCU activity, we treated human spermatozoa with the MCU inhibitor Ru360. This significantly prevented the increase in mitochondrial Ca^2+^ (11,080 ± 7,630) **(Figure 4H)**. To exclude the possibility that Ru360 indirectly reduced mitochondrial Ca^2+^ by interfering with CatSper-mediated Ca^2+^ entry, we assessed cytosolic Ca^2+^ responses using Fluo-4. CatSper was stimulated with progesterone, and Ca^2+^ responses were compared between control and Ru360-treated spermatozoa. Ru360 did not alter either basal cytosolic Ca^2+^ or the progesterone-induced Ca^2+^ response under capacitating conditions **(Supplementary Figure 3)**. Together, these findings support a conserved mechanism in mouse and human spermatozoa in which capacitation promotes CatSper-dependent Ca^2+^ entry followed by MCU-mediated mitochondrial Ca^2+^ uptake.

### CatSper and MCU activity contribute to mitochondrial activation and hyperactivation during capacitation

If CatSper-mediated Ca^2+^ entry and subsequent MCU-dependent mitochondrial Ca^2+^ uptake promote mitochondrial activation, then inhibition of either pathway should attenuate mitochondrial activation. To test this, spermatozoa were capacitated in the presence of vehicle, the CatSper inhibitor NNC 55-0396, or the MCU inhibitor Ru360 **(Figure 5A)**. The capacitation-associated increase in mitochondrial membrane potential was significantly reduced by inhibition of either CatSper or MCU (Ctrl: 8.66 ± 3.17; NNC: 6.25 ± 3.11; Ru360: 7.01 ± 3.11) **(Figure 5B)**. Thus, the capacitation-associated increase in mitochondrial membrane potential depends, at least in part, on CatSper and MCU activity.

**Figure 5.**
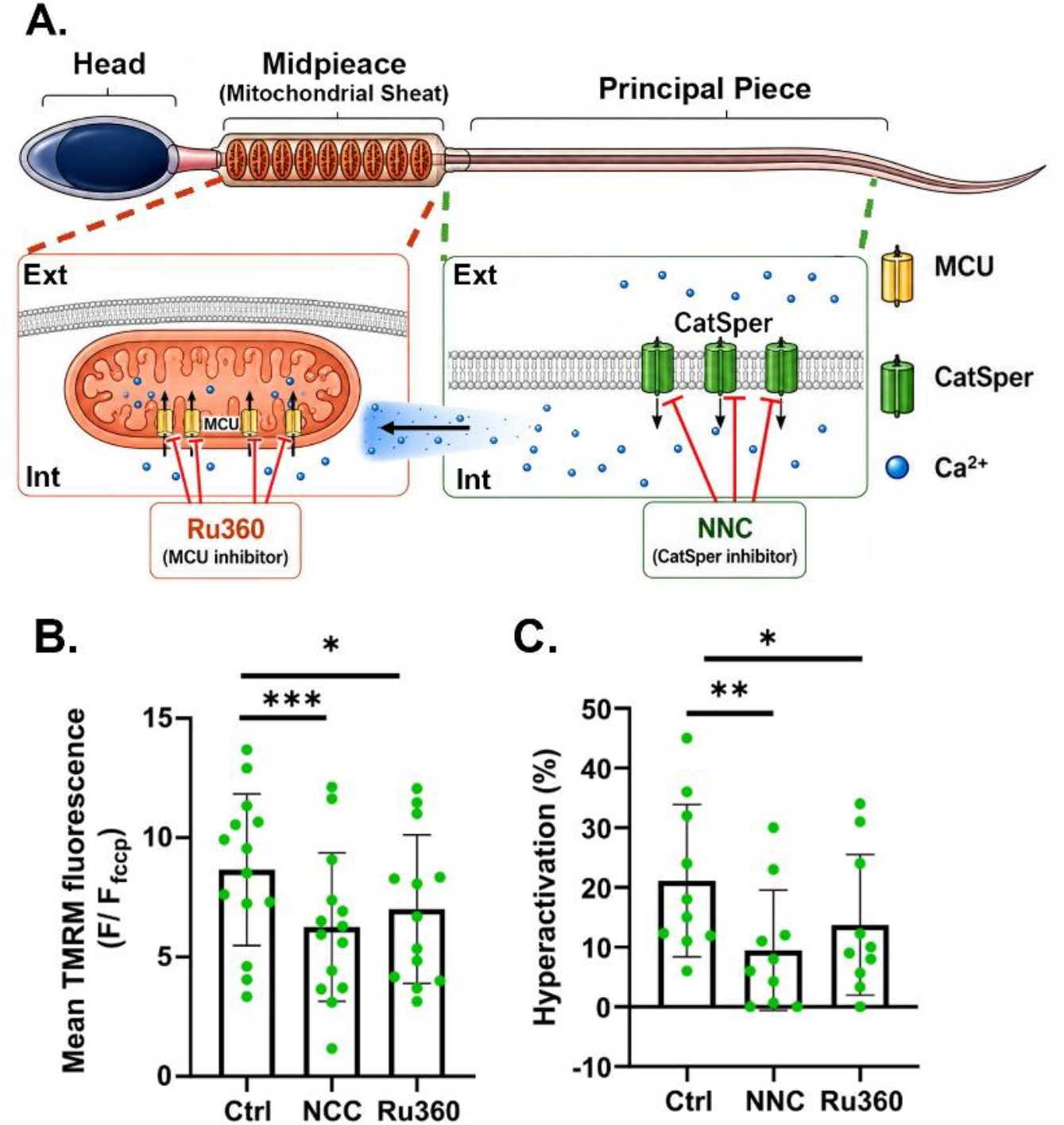
CatSper and MCU activity contribute to the capacitation-associated increase in mitochondrial membrane potential and hyperactivation. (**A**) Schematic representation of the localization and pharmacological inhibition of CatSper and the mitochondrial calcium uniporter (MCU) in human spermatozoa. CatSper is localized to the plasma membrane of the principal piece and is inhibited by NNC 55-0396 (NNC). MCU is localized to the inner mitochondrial membrane in the midpiece and is inhibited by Ru360. (**B–C**) Spermatozoa were capacitated for 4 h in the presence of vehicle (Ctrl), 1 µM NNC, or 20 µM Ru360. **(B)** Mitochondrial membrane potential assessed by flow cytometric analysis of TMRM fluorescence (n = 14). **(C)** Percentage of hyperactivated spermatozoa assessed by computer-assisted spermatozoa analysis (CASA) and expressed as a percentage of motile spermatozoa (n = 10). Flow cytometry and CASA measurements were performed at Washington University in St. Louis. * *P* < 0.05; ** *P* < 0.01; *** *P* < 0.001 by paired *t*-tests.

Finally, we asked whether disruption of this pathway also affected capacitation-associated hyperactivated motility, which increased from 3.45 ± 3.76% in NC spermatozoa to 21.12 ± 12.73% after capacitation. This increase was significantly attenuated when spermatozoa were capacitated in the presence of either NNC 55-0396 (9.50 ± 10.06%) or Ru360 (13.73 ± 11.77%) **(Figure 5C)**. Consistent with these findings, VCL after capacitation was also reduced following inhibition of either CatSper or MCU (Ctrl: 99.91 ± 19.15 µm/s; NNC: 79.82 ± 33.25 µm/s; Ru360: 82.82 ± 30.39 µm/s), whereas progressive motility was unaffected **(Supplementary Figure 4)**.

## Discussion

Together, our findings indicate that, as in mouse spermatozoa ^23^, CatSper-mediated Ca^2+^ entry promotes MCU-dependent mitochondrial Ca^2+^ uptake and mitochondrial activation, which in turn contributes to the acquisition of hyperactivated motility during human spermatozoa capacitation (**Figure 6**). This model is supported by the following principal findings. First, RCR, coupling efficiency, and mitochondrial membrane potential increase during human spermatozoa capacitation. Second, mitochondrial activity contributes to the acquisition of hyperactivated motility but is dispensable for the acrosome reaction. Third, the capacitation-associated increase in mitochondrial Ca^2+^ depends on extracellular Ca^2+^, CatSper activity, and MCU-mediated mitochondrial Ca^2+^ uptake. Fourth, inhibition of either CatSper or MCU attenuates the capacitation-associated increase in mitochondrial membrane potential and hyperactivation.

**Figure 6.**
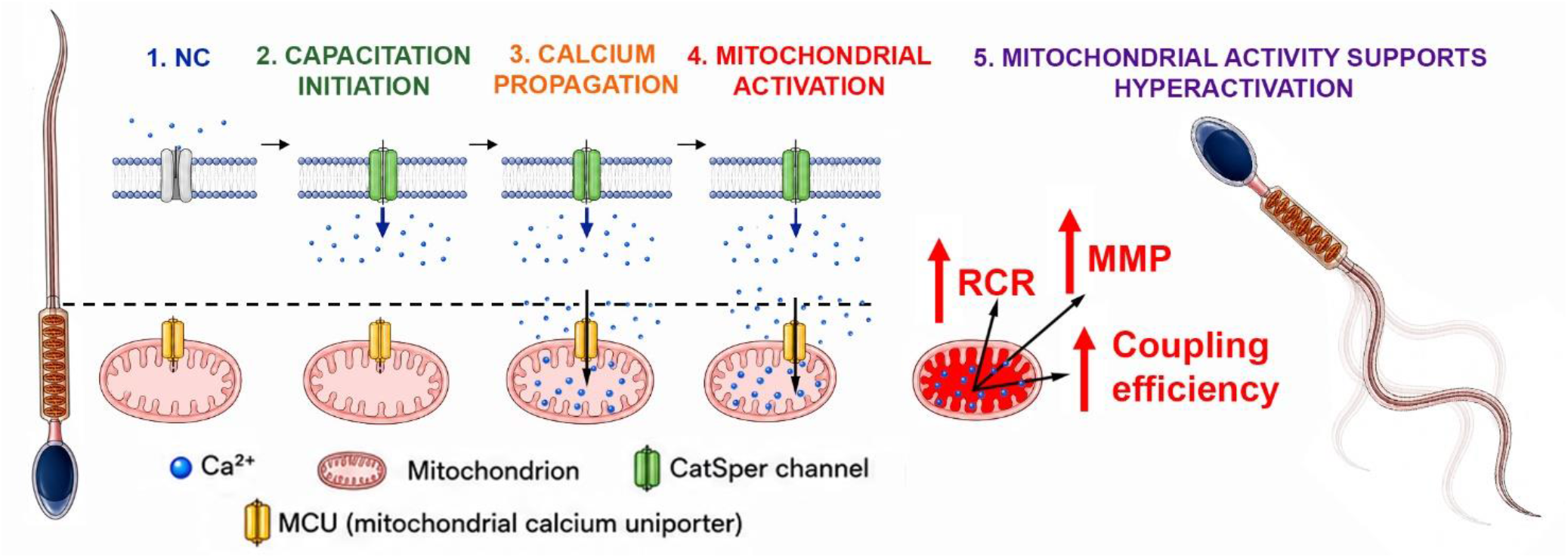
Proposed model linking CatSper-dependent Ca^2+^ signaling to mitochondrial activation and hyperactivated motility during human spermatozoa capacitation. (1) In non-capacitated (NC) spermatozoa, CatSper activity and cytosolic Ca^2+^ are low, resulting in limited mitochondrial Ca^2+^ uptake. (2) During capacitation, CatSper activation increases Ca^2+^ entry into the principal piece and elevates cytosolic Ca^2+^. (3) The cytosolic Ca^2+^ diffuses into the midpiece and is taken up into the mitochondrial matrix through the mitochondrial calcium uniporter (MCU), increasing mitochondrial Ca^2+^. (4) Increased mitochondrial Ca^2+^ stimulates Ca^2+^-sensitive mitochondrial enzymes and metabolic pathways, contributing to the capacitation-associated increases in mitochondrial membrane potential (MMP), respiratory control ratio (RCR), and coupling efficiency. (5) Enhanced mitochondrial activity contributes to the acquisition of hyperactivated motility during capacitation.

The role of mitochondria in mammalian spermatozoa function, bioenergetics, and fertilizing capacity has been debated for many years, in part because the relative contributions of glycolysis and oxidative phosphorylation vary among species and physiological conditions. In mouse spermatozoa, glycolytically generated ATP is sufficient to sustain progressive motility, whereas in human spermatozoa, mitochondrial function correlates strongly with spermatozoa motility and overall quality ^44,45^ . These observations have led to the concept that spermatozoa rely differently on glycolysis and mitochondrial metabolism depending on species and environmental conditions ^17,19,21,35,46,47^.

Increased mitochondrial activity during human spermatozoa capacitation has been previously reported, including by our group ^19,20,46^. Our findings are consistent with these previous observations and further support an increase in mitochondrial activity associated with capacitation.

Our finding that spare respiratory capacity did not increase despite increased mitochondrial respiration suggests that CAP human spermatozoa utilize a greater proportion of their available respiratory capacity than NC spermatozoa. This increased respiratory demand may be particularly relevant during the transition to hyperactivated motility, which requires increased energy ^48,49^. The accompanying decrease in NAD(P)H autofluorescence is consistent with an altered cellular redox state associated with increased respiratory activity. However, because NAD(P)H levels reflect the balance between production and oxidation, these measurements cannot distinguish between these processes.

The effects of mitochondrial disruption were not uniform across capacitation-associated functions. Treatment with AA or FCCP significantly reduced hyperactivation without producing a generalized loss of motility, whereas neither treatment affected spontaneous or Ca^2+^ ionophore-induced acrosome reaction. A similar effect has been observed in mouse spermatozoa ^23^, suggesting that the preferential contribution of mitochondrial activity to hyperactivated motility may be conserved across species. One possible explanation for this functional distinction is that hyperactivation and the acrosome reaction depend on different spatial and temporal patterns of Ca^2+^ signaling. Hyperactivation depends strongly on CatSper-mediated Ca^2+^ entry and flagellar Ca^2+^ signaling ^50^, whereas the acrosome reaction involves Ca^2+^ mobilization from intracellular stores together with Ca^2+^ entry across the plasma membrane ^51,52^. Distinct Ca^2+^ sources and spatially organized Ca^2+^ signals can therefore regulate different spermatozoa functions ^36,37^. The preservation of the acrosome reaction and overall motility despite impaired hyperactivation supports a preferential role for mitochondrial activity in the motility-related changes associated with capacitation.

In other cell types, entry of Ca^2+^ into mitochondria, primarily via MCU (De Stefani et al., 2011; Rossi et al., 2019a), plays an important role in both intracellular Ca^2+^ homeostasis and the regulation of oxidative metabolism ^25,54^. Following CatSper-mediated Ca^2+^ entry into the cytosol, Ca^2+^ could reach the mitochondrial intermembrane space through VDACs in the outer mitochondrial membrane, which are permeable to Ca^2+^ and may regulate its flux across the outer mitochondrial membrane ^55,56^. VDAC isoforms are also expressed in human spermatozoa ^57^, suggesting that this pathway could contribute to the transfer of Ca^2+^ to the mitochondrial compartment before its uptake into the matrix through MCU. Whether VDACs directly contribute to this process during sperm capacitation remains to be determined. In this context, we found that mitochondrial Ca^2+^ increased during human spermatozoa capacitation and that this increase was prevented by removal of extracellular Ca^2+^, inhibition of CatSper, or inhibition of MCU. These observations are consistent with our previous findings in mouse spermatozoa ^23^. Importantly, the MCU inhibitor Ru360 did not alter basal cytosolic Ca^2+^ or the Ca^2+^ response elicited by progesterone-induced CatSper activation, supporting the interpretation that its effect on mitochondrial Ca^2+^ was not due to inhibition of CatSper-mediated Ca^2+^ entry.

Inhibition of either CatSper or MCU also attenuated the capacitation-associated increase in mitochondrial membrane potential and reduced hyperactivated motility. CatSper is a well-established regulator of hyperactivation in mouse and human spermatozoa ^12,13,16,58^, and loss-of-function variants in human *CATSPER* genes impair hyperactivation despite otherwise relatively normal semen parameters ^16^. MCU-mediated Ca^2+^ uptake has previously been implicated in the maintenance of human sperm motility and ATP levels under conditions in which capacitation was not specifically assessed (Bravo et al., 2015). Together, these results place MCU downstream of CatSper-mediated Ca^2+^ entry and support a role for mitochondrial Ca^2+^ uptake in the development of hyperactivated motility during capacitation.

An important question is how mitochondrial Ca^2+^ uptake contributes to hyperactivation. One possibility is that mitochondrial Ca^2+^ stimulates oxidative metabolism to help meet the increased energetic demands of hyperactivated motility. In somatic cells, mitochondrial Ca^2+^ regulates several enzymes involved in oxidative metabolism ^22,25,53^. Consistent with this possibility, we found that capacitation increased mitochondrial Ca^2+^, mitochondrial membrane potential, and coupling efficiency, whereas inhibition of MCU reduced hyperactivation. Previous work in ejaculated human spermatozoa showed that capacitation increases both glycolysis and oxidative phosphorylation and that both pathways contribute to hyperactivation ^60^. However, because we did not directly measure ATP production or glycolytic flux, our findings do not establish whether mitochondrial Ca^2+^ uptake promotes hyperactivation by increasing ATP production or through another mitochondrial mechanism.

A second, non-exclusive possibility is that mitochondrial Ca^2+^ uptake contributes to the spatial and temporal regulation of cytosolic Ca^2+^ signals. CatSper channels are localized to the principal piece, whereas the mitochondrial sheath is located in the midpiece. Given this spatial separation, CatSper-dependent Ca^2+^ signals reaching the midpiece and subsequent mitochondrial Ca^2+^ uptake could influence the magnitude, duration, or spatial distribution of cytosolic Ca^2+^ signals and thereby modulate flagellar behavior. Whether mitochondrial Ca^2+^ promotes hyperactivation primarily through regulation of mitochondrial metabolism, modulation of Ca^2+^ signaling, or a combination of these mechanisms remains to be determined.

Several limitations should be considered when interpreting these findings. Most importantly, the inhibitors used in this study are not completely selective. NNC 55-0396 can inhibit the sperm potassium channel SLO3 at concentrations similar to those used to inhibit CatSper ^61^ and can affect several human sperm motility parameters and the acrosome reaction ^62,63^. Ru360 can also reduce total motility and ATP content in human spermatozoa ^59^. Under our experimental conditions, however, neither NNC 55-0396 nor Ru360 altered total or progressive motility despite reducing hyperactivation, arguing against a generalized impairment of sperm motility. Nevertheless, potential off-target effects of these inhibitors cannot be fully excluded. An additional limitation is that we used CASA to evaluate spermatozoa motility at the population level. Although CASA provides robust quantification of conventional motility parameters and hyperactivation, it cannot capture subtle alterations in flagellar dynamics or specific swimming behaviors. Thus, future studies should use higher-resolution approaches to examine individual flagellar dynamics and swimming patterns. Finally, whether mitochondrial Ca^2+^ uptake is required for fertilization in human spermatozoa remains unknown. For ethical reasons, the effects of mitochondrial or MCU inhibition on human fertilization cannot be directly tested as they can in animal models. Additional approaches will therefore be required to further define the molecular mechanisms involved and their relevance to fertilization.

In conclusion, our findings support a model in which CatSper-mediated Ca^2+^ entry promotes MCU-dependent mitochondrial Ca^2+^ uptake and mitochondrial activation, thereby contributing to the acquisition of hyperactivated motility during human sperm capacitation. These findings identify mitochondrial Ca^2+^ handling as an important component of the signaling network coupling capacitation-associated Ca^2+^ entry to changes in sperm motility.

## Contributor Information

Celia M. Santi.

Rossana Sapiro.

## Data Availability Statement

The raw data supporting the conclusions of this article will be made available by the authors, without undue reservation.

## Author Contributions

PI and JF conceived and designed the study, performed experiments, analyzed the data, and interpreted the results. SB and NP performed experiments and analyzed the data. CMS, AC, and RS contributed to data analysis and interpretation of the results. JF and PI prepared the figures and drafted the manuscript. PI, JF, AC, SB, NP, CMS, and RS critically reviewed and revised the manuscript. All authors contributed to the article and read and approved of the final submitted version.

## Acknowledgments

We thank Dr. Deborah Frank, Scientific Editor in the Department of Obstetrics and Gynecology at Washington University School of Medicine, for her critical review of the manuscript. We also thank the Fertility and Reproductive Medicine Center at Washington University School of Medicine in St. Louis and the Fertilab-Reprovita andrology clinics in Montevideo, Uruguay, for providing human semen samples used in this study.

## References

1. Yanagimachi, R. The Movement Of Golden Hamster Spermatozoa Before And After Capacitation. Reproduction 23, 193–196 (1970).

2. Demott, R. P. & Suarez, S. S. Hyperactivated Sperm Progress in the Mouse Oviduct1. Biol. Reprod. 46, 779–785 (1992).

3. Suarez, S. S., Varosi, S. M. & Dai, X. Intracellular calcium increases with hyperactivation in intact, moving hamster sperm and oscillates with the flagellar beat cycle. Proceedings of the National Academy of Sciences 90, 4660–4664 (1993).

4. Stival, C. et al. Sperm Capacitation and Acrosome Reaction in Mammalian Sperm. in 93–106 (2016). doi:10.1007/978-3-319-30567-7_5.

5. Hirohashi, N. & Yanagimachi, R. Sperm acrosome reaction: its site and role in fertilization†. Biol. Reprod. 99, 127–133 (2018).

6. Mariano G. Buffone. Sperm Acrosome Biogenesis and Function During Fertilization. vol. 220 (Springer International Publishing, Cham, 2016).

7. Puga Molina, L. C. et al. Molecular Basis of Human Sperm Capacitation. Front. Cell Dev. Biol. 6, (2018).

8. Carrasquel Martínez, G., Aldana, A., Carneiro, J., Treviño, C. L. & Darszon, A. Acrosomal alkalinization occurs during human sperm capacitation. Mol. Hum. Reprod. 28, (2022).

9. Lishko, P. V & Mannowetz, N. CatSper: a unique calcium channel of the sperm flagellum. Curr. Opin. Physiol. 2, 109–113 (2018).

10. Rahban, R. & Nef S. CatSper: The complex main gate of calcium entry in mammalian spermatozoa. Mol. Cell. Endocrinol. 518, 110951 (2020).

11. Ren, D. et al. A sperm ion channel required for sperm motility and male fertility. Nature 413, 603–609 (2001).

12. Carlson, A. E. et al. CatSper1 required for evoked Ca2+ entry and control of flagellar function in sperm. Proceedings of the National Academy of Sciences 100, 14864–14868 (2003).

13. Quill, T. A. et al. Hyperactivated sperm motility driven by CatSper2 is required for fertilization. Proceedings of the National Academy of Sciences 100, 14869–14874 (2003).

14. Vyklicka, L. & Lishko, P. V. Dissecting the signaling pathways involved in the function of sperm flagellum. Curr. Opin. Cell Biol. 63, 154–161 (2020).

15. Xia, J., Reigada, D., Mitchell, C. H. & Ren, D. CATSPER Channel-Mediated Ca2+ Entry into Mouse Sperm Triggers a Tail-to-Head Propagation1. Biol. Reprod. 77, 551–559 (2007).

16. Young, S. et al. Human fertilization in vivo and in vitro requires the CatSper channel to initiate sperm hyperactivation. Journal of Clinical Investigation 134, (2024).

17. Barbagallo, F. et al. Evaluation of Sperm Mitochondrial Function: A Key Organelle for Sperm Motility. J. Clin. Med. 9, 363 (2020).

18. Giaccagli, M. M. et al. Capacitation-Induced Mitochondrial Activity Is Required for Sperm Fertilizing Ability in Mice by Modulating Hyperactivation. Front. Cell Dev. Biol. 9, (2021).

19. Carrageta, D. F. et al. Mitochondrial Activation and Reactive Oxygen-Species Overproduction during Sperm Capacitation are Independent of Glucose Stimuli. Antioxidants 9, 750 (2020).

20. Irigoyen, P., Mansilla, S., Castro, L., Cassina, A. & Sapiro, R. Mitochondrial function and reactive oxygen species production during human sperm capacitation: Unraveling key players. The FASEB Journal 38, (2024).

21. Kumar, N. Sperm Mitochondria, the Driving Force Behind Human Spermatozoa Activities: Its Functions and Dysfunctions-A Narrative Review. Curr. Mol. Med. 23, 332–340 (2023).

22. Rossi, A., Pizzo, P. & Filadi R. Calcium, mitochondria and cell metabolism: A functional triangle in bioenergetics. Biochimica et Biophysica Acta (BBA)-Molecular Cell Research 1866, 1068–1078 (2019).

23. Ferreira, J. J. et al. Increased mitochondrial activity upon CatSper channel activation is required for mouse sperm capacitation. Redox Biol. 48, 102176 (2021).

24. Rossi, A., Pizzo, P. & Filadi, R. Calcium, mitochondria and cell metabolism: A functional triangle in bioenergetics. Biochimica et Biophysica Acta (BBA)-Molecular Cell Research 1866, 1068–1078 (2019).

25. Denton, R. M. Regulation of mitochondrial dehydrogenases by calcium ions. Biochimica et Biophysica Acta (BBA)-Bioenergetics 1787, 1309–1316 (2009).

26. Kamer, K. J. & Mootha, V. K. The molecular era of the mitochondrial calcium uniporter. Nat. Rev. Mol. Cell Biol. 16, 545–553 (2015).

27. Ferreira, J. J., Lybaert, P., Puga-Molina, L. C. & Santi, C. M. Conserved Mechanism of Bicarbonate-Induced Sensitization of CatSper Channels in Human and Mouse Sperm. Front. Cell Dev. Biol. 9, (2021).

28. Brenker, C. et al. The CatSper channel: a polymodal chemosensor in human sperm. EMBO J. 31, 1654–1665 (2012).

29. Jeschke, J. K. et al. The Action of Reproductive Fluids and Contained Steroids, Prostaglandins, and Zn2+ on CatSper Ca2+ Channels in Human Sperm. Front. Cell Dev. Biol. 9, (2021).

30. Edwards, M. et al. ABHD2 activity is not required for the non-genomic action of progesterone on human sperm. Human Reproduction 41, 1409–1419 (2026).

31. Strünker, T. et al. The CatSper channel mediates progesterone-induced Ca2+ influx in human sperm. Nature 471, 382–386 (2011).

32. Mannowetz, N., Miller, M. R. & Lishko, P. V. Regulation of the sperm calcium channel CatSper by endogenous steroids and plant triterpenoids. Proceedings of the National Academy of Sciences 114, 5743–5748 (2017).

33. Lishko, P. V & Mannowetz, N. CatSper: a unique calcium channel of the sperm flagellum. Curr. Opin. Physiol. 2, 109–113 (2018).

34. Miller, M. R. et al. Unconventional endocannabinoid signaling governs sperm activation via the sex hormone progesterone. Science (1979). 352, 555–559 (2016).

35. Bravo, A. et al. Disruption of Calcium Homeostasis in Human Spermatozoa: Implications on Mitochondrial Bioenergetics, ROS Production, Phosphatidylserine Externalization, and Motility. Antioxidants 15, 213 (2026).

36. Alasmari, W. et al. The clinical significance of calcium-signalling pathways mediating human sperm hyperactivation. Human Reproduction 28, 866–876 (2013).

37. Alasmari, W. et al. Ca2+ Signals Generated by CatSper and Ca2+ Stores Regulate Different Behaviors in Human Sperm*. Journal of Biological Chemistry 288, 6248–6258 (2013).

38. World-Health-Organization, 2010. 5th ed ed. Geneva. s.l.:World Health Organization.. WHO Laboratory Manual for the Examination and Processing of Human Semen. . (2021).

39. Björndahl, L. et al. Standards in semen examination: publishing reproducible and reliable data based on high-quality methodology. Human Reproduction 37, 2497–2502 (2022).

40. Marín-Briggiler, C. I. et al. Human Sperm Remain Motile After a Temporary Energy Restriction but do Not Undergo Capacitation-Related Events. Front. Cell Dev. Biol. 9, (2021).

41. Mortimer, D. & Mortimer, S. T. Computer-Aided Sperm Analysis (CASA) of Sperm Motility and Hyperactivation. in 77–87 (2013). doi:10.1007/978-1-62703-038-0_8.

42. Irigoyen, P., Sapiro, R. & Cassina, A. High-Resolution Respirometry to Assess Mitochondrial Function in Human Spermatozoa. Journal of Visualized Experiments https://doi.org/10.3791/65493 (2023) doi:10.3791/65493.

43. Brand, M. D. & Nicholls, D. G. Assessing mitochondrial dysfunction in cells. Biochemical Journal 435, 297–312 (2011).

44. Irigoyen, P. et al. Mitochondrial metabolism determines the functional status of human sperm and correlates with semen parameters. Front. Cell Dev. Biol. 10, (2022).

45. Cassina, A. et al. Defective Human Sperm Cells Are Associated with Mitochondrial Dysfunction and Oxidant Production1. Biol. Reprod. 93, (2015).

46. Stendardi, A. et al. Evaluation of mitochondrial respiratory efficiency during in vitro capacitation of human spermatozoa. Int. J. Androl. 34, 247–255 (2011).

47. Tourmente, M., Sansegundo, E., Rial, E. & Roldan, E. R. S. Capacitation promotes a shift in energy metabolism in murine sperm. Front. Cell Dev. Biol. 10, (2022).

48. Ho, H.-C., Granish, K. A. & Suarez, S. S. Hyperactivated Motility of Bull Sperm Is Triggered at the Axoneme by Ca2+ and Not cAMP. Dev. Biol. 250, 208–217 (2002).

49. Sansegundo, E., Tourmente, M. & Roldan, E. R. S. Energy Metabolism and Hyperactivation of Spermatozoa from Three Mouse Species under Capacitating Conditions. Cells 11, 220 (2022).

50. Hwang, J. Y. & Chung, J.-J. CatSper Calcium Channels: 20 Years On. Physiology 38, 125–140 (2023).

51. Costello, S. et al. Ca2+-stores in sperm: their identities and functions. Reproduction 138, 425–437 (2009).

52. Correia, J., Michelangeli, F. & Publicover, S. Regulation and roles of Ca2+ stores in human sperm. Reproduction 150, R65–R76 (2015).

53. De Stefani, D., Raffaello, A., Teardo, E., Szabò, I. & Rizzuto, R. A forty-kilodalton protein of the inner membrane is the mitochondrial calcium uniporter. Nature 476, 336–340 (2011).

54. Contreras, L., Drago, I., Zampese, E. & Pozzan, T. Mitochondria: The calcium connection. Biochimica et Biophysica Acta (BBA)-Bioenergetics 1797, 607–618 (2010).

55. Rosencrans, W. M., Rajendran, M., Bezrukov, S. M. & Rostovtseva, T. K. VDAC regulation of mitochondrial calcium flux: From channel biophysics to disease. Cell Calcium 94, 102356 (2021).

56. Sander, P., Gudermann, T. & Schredelseker, J. A Calcium Guard in the Outer Membrane: Is VDAC a Regulated Gatekeeper of Mitochondrial Calcium Uptake? Int. J. Mol. Sci. 22, 946 (2021).

57. Liu, B., Wang, Z., Zhang, W. & Wang, X. Expression and localization of voltage-dependent anion channels (VDAC) in human spermatozoa. Biochem. Biophys. Res. Commun. 378, 366–370 (2009).

58. Ren, D. et al. A sperm ion channel required for sperm motility and male fertility. Nature 413, 603–609 (2001).

59. Bravo, A. et al. Effect of mitochondrial calcium uniporter blocking on human spermatozoa. Andrologia 47, 662–668 (2015).

60. Balbach, M. et al. Capacitation induces changes in metabolic pathways supporting motility of epididymal and ejaculated sperm. Front. Cell Dev. Biol. 11, (2023).

61. Rennhack, A. et al. A novel cross-species inhibitor to study the function of CatSper Ca2+ channels in sperm. Br. J. Pharmacol. 175, 3144–3161 (2018).

62. Tamburrino, L. et al. The CatSper calcium channel in human sperm: relation with motility and involvement in progesterone-induced acrosome reaction. Human Reproduction 29, 418–428 (2014).

63. Maleki, S. et al. Effects of CatSper Stimulation and Inhibition by Progesterone and NNC on Human Sperm. Andrologia 2023, (2023).

