## Supplementary Figures for "Mitochondrial Calcium Uptake Promotes Mitochondrial Activation and Hyperactivated Motility During Human Sperm Capacitation"

**Supplementary material.**

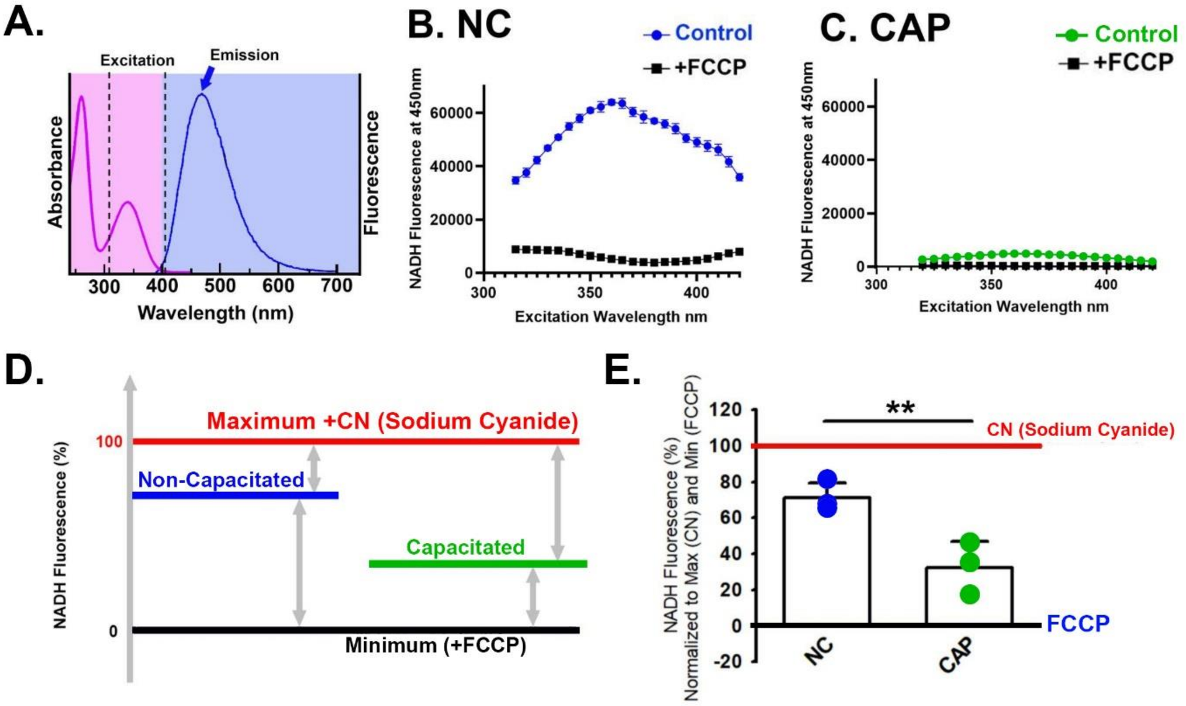


**Supplementary Figure 1. Validation and quantification of NADH autofluorescence in non-capacitated and capacitated human spermatozoa.** (**A**) Excitation and emission wavelengths used to measure NADH autofluorescence. (**B–C**) Excitation spectra of NADH autofluorescence recorded at an emission wavelength of 450 nm in non-capacitated (NC; blue, **B**) and capacitated (CAP; green, **C**) spermatozoa under control conditions and after FCCP treatment (black). These spectra were used to validate the excitation wavelength selected for subsequent NADH measurements. **(D)** Schematic representation of the fluorescence normalization approach using FCCP-induced minimum fluorescence and sodium cyanide (CN)-induced maximum fluorescence as reference points. **(E)** Quantification of NADH autofluorescence (λ_ex = 310 nm; λ_em = 450 nm) following normalization between the FCCP-defined minimum and CN-defined maximum. NC and CAP samples were obtained from the same donor and analyzed in parallel following the capacitation period. **P* < 0.05; ***P* < 0.01; ****P* < 0.001 by paired *t*-test.

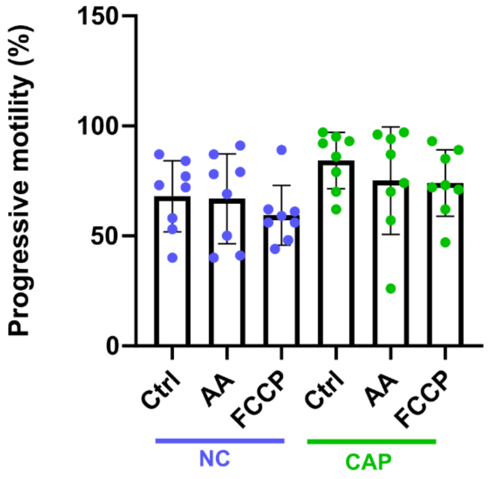


**Supplementary Figure 2. Effects of mitochondrial disruption on sperm progressive motility (a+b,%).** Spermatozoa motility was assessed by CASA at the Facultad de Medicina, Universidad de la República (UDELAR). **M**otility parameters were evaluated in the presence of the mitochondrial disruptors antimycin A (AA) and FCCP. *n* = 8. NC, non-capacitated (blue dots); CAP, capacitated (green dots). No significant differences were observed.

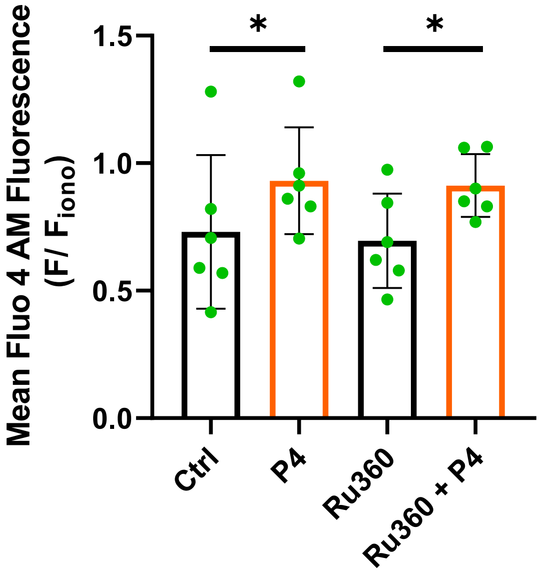


**Supplementary Figure 3. Ru360 does not affect the progesterone-induced increase in intracellular $Ca^{2+}$ in human spermatozoa.** Intracellular $Ca^{2+}$ was measured using Fluo-4 by flow cytometry in capacitated (CAP) spermatozoa in the absence (Ctrl) or presence of the MCU inhibitor Ru360 ($n$ = 6) with or without Progesterone (P4; 30 µM). Data were acquired by flow cytometry at Washington University in St. Louis. *$P < 0.05$ by paired t -test.

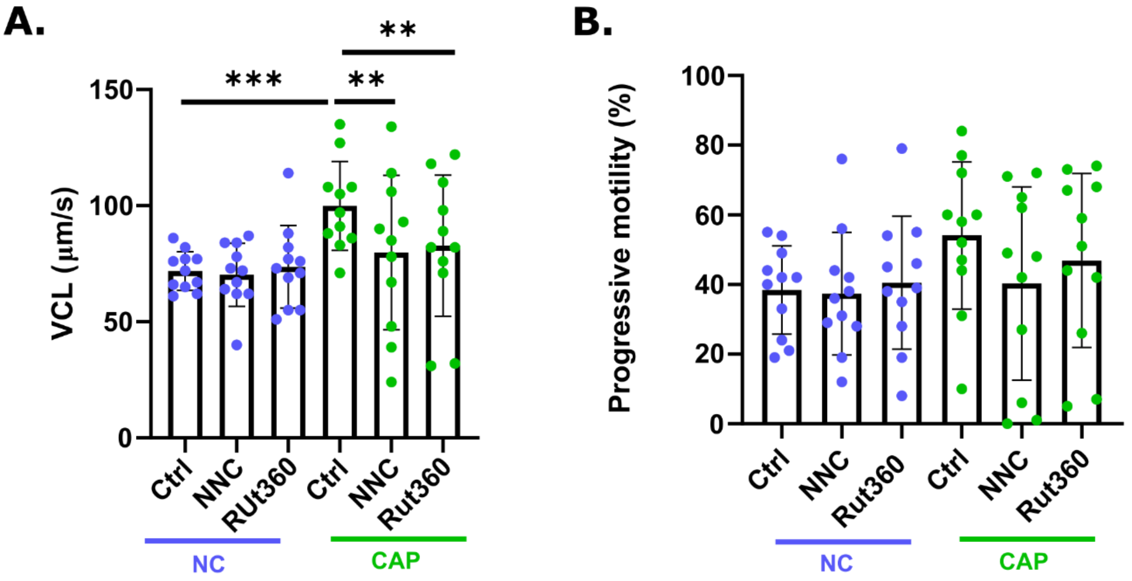


**Supplementary Figure 4. Effects of $Ca^{2+}$ channel inhibition on spermatozoa motility parameters.** Spermatozoa motility was assessed by CASA at Washington University in St. Louis (WashU) Motility parameters were evaluated in the presence of the CatSper inhibitor NNC 55-0396 (NNC) and the MCU inhibitor Ru360. **(A)** Curvilinear velocity (VCL), n=11. **(B)** Progressive motility (a+b, %), $n$ = 11. NC, non-capacitated (blue dots); CAP, capacitated (green dots). $^{**}P < 0.01$; $^{***}P < 0.001$ by paired t-test.
